# Identification of TAZ as the essential molecular switch in orchestrating SCLC phenotypic transition and metastasis

**DOI:** 10.1101/2021.07.28.454244

**Authors:** Yujuan Jin, Qiqi Zhao, Weikang Zhu, Yan Feng, Tian Xiao, Peng Zhang, Liyan Jiang, Yingyong Hou, Chenchen Guo, Hsinyi Huang, Yabin Chen, Xinyuan Tong, Jiayu Cao, Fei Li, Xueliang Zhu, Jun Qin, Dong Gao, Xin-Yuan Liu, Hua Zhang, Luonan Chen, Roman K Thomas, Kwok-Kin Wong, Yong Wang, Liang Hu, Hongbin Ji

## Abstract

Small cell lung cancer (SCLC) is a recalcitrant cancer featured with high metastasis. However, the exact cell type contributing to metastasis remains elusive. Using *Rb1*^L/L^*/Trp53*^L/L^ mouse model, we identify the NCAM^hi^CD44^lo/−^ subpopulation as SCLC metastasizing cell (SMC), which is progressively transitioned from non-metastasizing NCAM^lo^CD44^hi^ cell (Non-SMC). Integrative chromatin accessibility and gene expression profiling studies reveal an important role of SWI/SNF complex, and knockout of its central component, *Brg1*, significantly inhibits such phenotypic transition and metastasis. Mechanistically, TAZ is silenced by SWI/SNF complex during SCLC malignant progression, and its knockdown promotes SMC transition and metastasis. Importantly, ectopic TAZ expression reversely drives SMC-to-Non-SMC transition and alleviates metastasis. Single-cell RNA-sequencing analyses identify SMC as the dominant subpopulation in human SCLC metastasis, and immunostaining data show a positive correlation between TAZ and patient prognosis. These data uncover high SCLC plasticity and identify TAZ as key molecular switch in orchestrating SCLC phenotypic transition and metastasis.

## Introduction

Small cell lung cancer (SCLC) features with very poor prognosis, with about 15% global lung cancer incidence and a five-year survival lower than 7% ^1^. This can be largely attributed to the highly metastatic capability of SCLC. Most SCLC patients are initially diagnosed at extensive stage, characterized with nearby lung and/or distant metastases. Therefore, it’s urgently needed to explore the mechanisms involved in SCLC metastasis so as to provide helpful insights into clinical management.

Previous studies have shown that approximately 90% of human SCLC harbor concurrent inactivating mutations or deletions of *Rb1* and *Trp53* ^2^. Homozygous deletion of these two alleles in mouse lung epithelia promotes SCLC development and dramatic metastasis, which closely recapitulates human SCLC in the clinic ^3^. Mouse SCLC in *Rb1*^L/L^/*Trp53*^L/L^ (RP) model typically express neuroendocrine markers including neuronal cell adhesion molecule (NCAM) and achaete-scute complex homolog 1 (ASCL1), and frequently metastasize into distant organs ^3^. Concurrent deletion of *P130*, an *Rb*-related gene, or *Pten* in RP model significantly accelerates malignant progression and SCLC metastasis ^4, 5^. Moreover, up-regulated NFIB expression is found to promote SCLC metastasis through increasing the accessibility of global chromatin ^6–8^.

SCLC are featured with high heterogeneity ^5–15^. It is proposed that human SCLC is composed of four different subtypes based on lineage-related transcription factors including ASCL1, NEUROD1, YAP, and POU2F3 ^14^. More recently, an inflamed SCLC subtype has been identified with a good response to immunotherapy ^16^. Similar heterogeneity has also been found in mouse SCLC, e.g., the CD24^hi^CD44^lo^EpCAM^hi^ subpopulation from *RP* model is identified to harbor strong capability to form tumors in allograft assay ^11^. Moreover, mouse SCLC are found to contain the neuroendocrine (NE) and non-neuroendocrine (non-NE) subpopulations according to distinct growth patterns in culture, with the NE subtype growing as suspension and the non-NE as adhesion ^9^. The NE cells frequently express neuroendocrine markers including NCAM, synaptophysin (SYP) and ASCL1. In contrast, the non-NE cells tend to express mesenchymal markers such as VIMENTIN and CD44 ^9^. It has been reported that the synergetic cooperation between NE and non-NE subpopulations is necessary for SCLC metastasis whereas neither subtype could metastasize on its own ^9^. Therefore, the exact population responsible for SCLC metastasis still remains unknown.

The Switch/Sucrose-Nonfermentable (mSWI/SNF) complexes, including canonical BRG1/BRM-associated factor (BAF), polybromo-associated BAF (PBAF) and non-canonical BAF (ncBAF), are essential for chromatin remodeling ^17 18^. All the three complexes contain a core ATPase subunit, e.g., BRG1 (Brahma/SWI2-related gene 1, also called SMARCA4), which catalyzes the hydrolysis of ATP ^18^. Previous studies reveal that the SWI/SNF complexes tend to function as the tumor suppressor during cancer development. Consistently, a high incidence of BRG1 inactivating mutation is detected in multiple cancer types including lung cancer ^19^. Previous studies show that BRG1 promotes cell cycle arrest and senescence through the retinoblastoma pathway in cancer cells ^20, 21^. Interestingly, recent studies have also indicated an oncogenic role of BRG1. For example, BRG1 promotes pancreatic intraepithelial neoplasia (PanIN) development and gastric cancer metastasis ^22, 23, 24^. In SCLC, BRG1 is preferentially required for cancer progression when MAX (Myc-associated factor) is inactivated ^25^. These findings indicate that BRG1 might function as tumor suppressor or oncogenic driver in cell type- or genetic-context dependent manner.

The Hippo pathway is initially defined as an important pathway during organ size control and function mainly through the synergetic interaction between the transcription factors TEAD1-4 and transcriptional co-activator YAP/TAZ (WWTR1) ^26^. The oncogenic activities of YAP/TAZ have been well documented in multiple epithelial cancers ^26–31^. It is well-known that YAP/TAZ sustain self-renewal and tumor-initiating capability, and promote cancer malignant progression and metastasis through epithelial to mesenchymal transition (EMT) ^30^. Latest studies also reveal that YAP/TAZ might function as a tumor suppressor ^32–34^. For instance, YAP expression is down-regulated in breast cancer and knockdown of YAP promotes cancer cells migration and invasiveness ^35^. Moreover, we have previously found YAP acts as the barrier for adenocarcinoma to squamous carcinoma transdifferetiation (AST) as well as lung squamous cell carcinoma progression ^33, 36^. Nonetheless, the exact role of YAP/TAZ during SCLC metastasis has not been characterized yet.

We here identify the NCAM^hi^CD44^lo/−^ cells as the major subpopulation responsible for SCLC metastasis. Moreover, this subpopulation is progressively transitioned from the non-metastatic NCAM^lo^CD44^hi^ cells via the SWI/SNF complex-mediated TAZ silencing. Our data highlight the important link between epigenetically regulated TAZ and SCLC plasticity and metastasis.

## Results

### Identification of the NCAM^hi^CD44^lo^/-subpopulation as SCLC metastasizing cells

To study SCLC heterogeneity in link with cancer malignant progression and metastasis, we first performed immunohistochemistry (IHC) staining in RP tumors using neuroendocrine marker NCAM and mesenchymal marker CD44. In primary RP tumors, we indeed observed a heterogeneous expression pattern of these two markers (**Fig. 1A**). We found that the percentage of NCAM^hi^CD44^lo/−^ tumors, defined with over 50% of cancer cells highly expressing NCAM and with low or no CD44 expression ^37^, increased with malignant progression and metastasis (**Fig. 1B and Fig. S1A, Table S1**). Consistently, we found that distant organ metastases such as liver and kidney metastases uniformly exhibited the NCAM^hi^CD44^lo/−^ expression pattern (**Fig. 1A**). These data indicate that the NCAM^hi^CD44^lo/−^ subpopulation might be responsible for SCLC metastasis.

**Figure 1.**
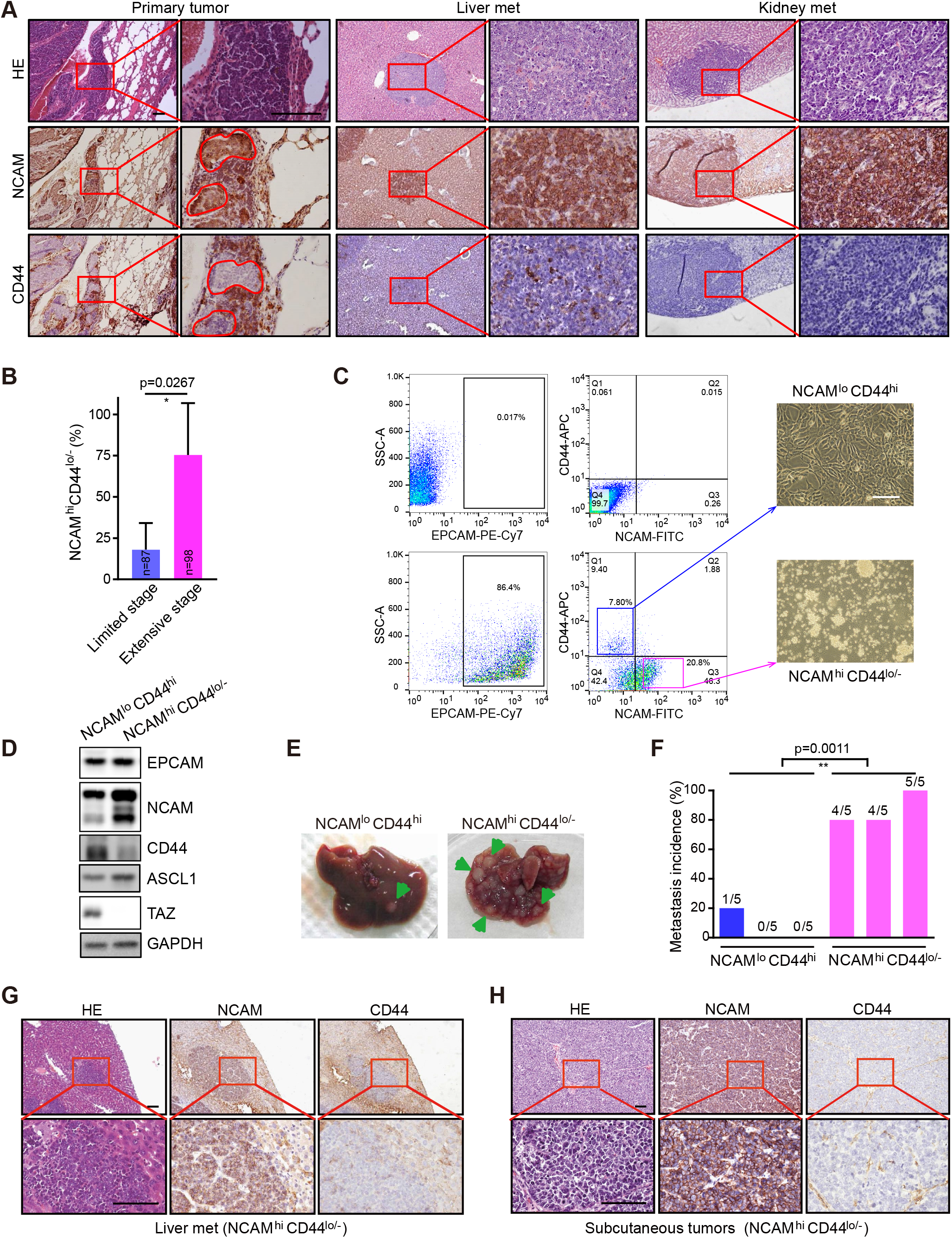
Identification of the NCAM^hi^CD44^lo/−^ cells as SCLC metastaizing cells in RP mouse model. **(A)** Representative photos of HE staining, NCAM and CD44 IHC staining of primary tumors, liver and kidney metastases (met) from RP mouse model. Scale bars, 100 μm. **(B)** Statistic analyses of the NCAM^hi^CD44^lo/−^ tumors at limited stage (no overt distant organ metastasis) and extensive stage (overt metastasis) in RP model. The NCAM^hi^CD44^lo/−^ tumors were defined when the lesions contained more than 50% cells showing NCAM^hi^ and CD44^lo/−^ expression. Limited stage: 87 tumors from 4 mice were analyzed; extensive stage: 98 tumors from 4 mice were analyzed. Data were shown as mean ± S.E.M. P value was calculated by unpaired two-tailed *t test*. **(C)** Flow Cytometry (FACS) analyses of primary tumors from RP mouse model using antibodies towards EPCAM, NCAM and CD44. The tumors cells without primary antibody incubation were showed as negative control (Top panels). The NCAM^lo^CD44^hi^ and NCAM^hi^CD44^lo/−^ cells were sorted and cultured *in vitro* and the representative cell growth photos were shown on the right. Scale bar, 100 μm. **(D)** Western blot detection of EPCAM, NCAM, CD44, ASCL1 and TAZ expression in established NCAM^lo^CD44^hi^ and NCAM^hi^CD44^lo/−^ SCLC primary cell lines. **(E-F)** Representative photos (**E**) and the incidence (**F**) of liver metastasis in nude mice subcutaneously transplanted with primary NCAM^lo^CD44^hi^ or NCAM^hi^CD44^lo/−^ cells derived from RP mouse model. Data were shown from three independent experiments (n=5 mice for each experiment). The ratio of mice with liver metastasis was also indicated. P value was calculated by unpaired two-tailed *t test*. **(G-H)** Representative photos of HE staining, NCAM and CD44 IHC staining of liver metastases (**G**), and subcutaneous tumors (**H**) in nude mice transplanted with NCAM^hi^CD44^lo/−^ cell lines. Scale bars, 100 μm.

To test this, we then used FACS sorting to isolate the NCAM^hi^CD44^lo/−^ and NCAM^lo^CD44^hi^ subpopulations from primary RP tumors (**Fig. 1C**). Genotyping analyses confirmed the concurrent deletion of *Rb1* and *Trp53* in both subpopulations (**Fig. S1B**). We found that the NCAM^hi^CD44^lo/−^ cells grew in culture as oncosphere with suspension growth pattern (**Fig. 1C**), similar to classical human SCLC cell lines. In contrast, the NCAM^lo^CD44^hi^ cells grew as adhesion (**Fig. 1C**). Moreover, higher ASCL1 level was detected in the NCAM^hi^CD44^lo/−^ subpopulation (**Fig. 1D**). We then subcutaneously transplanted 5 × 10^6^ cells from either NCAM^hi^CD44^lo/−^ or NCAM^lo^CD44^hi^ subpopulation into nude mice and waited for up to 10 weeks for distant organ metastasis analyses. Both subpopulations formed subcutaneous tumors at 100% percentage in allograft assay (**Fig. S1C**). In contrast, the metastasis analyses revealed a huge difference. Most mice (13 out of 15) from the NCAM^hi^CD44^lo/−^ group had spontaneous metastases in liver whereas only 1 out of 15 mice from the NCAM^lo^CD44^hi^ group displayed distant metastasis (**Fig. 1E-F**). The liver metastases from the NCAM^hi^CD44^lo/−^ group exhibited characteristic marker expression pattern, similar to subcutaneous tumors (**Fig. 1G-H**). These data demonstrate that the NCAM^hi^CD44^lo/−^ cells are mainly responsible for SCLC metastasis. We hereafter referred to the NCAM^hi^CD44^lo/−^ and NCAM^lo^CD44^hi^ subpopulations as SCLC metastasizing cell (SMC) and Non-SCLC metastasizing cell (Non-SMC), respectively.

### Phenotypic transition from Non-SMC to SMC contributes to SCLC metastasis

Consistent with SMC metastases tumors, the liver metastasis lesion from Non-SMC allograft assay also exhibited the NCAM^hi^CD44^lo/−^ expression pattern (**Fig. 2A and Fig. S2A**). We speculated that there might exist phenotypic transition from Non-SMC to SMC during SCLC malignant progression. To test this, we established Non-SMC cell line stably expressing GFP, Non-SMC-GFP, and performed subcutaneous allograft assay (**Fig. 2B**). Immunofluorescence (IF) staining in allograft tumors revealed that about 13±2% GFP-positive cancer cells displayed the NCAM^hi^CD44^lo/−^ pattern whereas the rest remained as Non-SMC expression pattern (**Fig. 2C and Table S2**). Consistently, both suspension and adhesion growth patterns were observed when these allograft tumors were cultured *in vitro* (**Fig. 2B**). To further confirm such transition, we picked single-cell clones from Non-SMC-GFP cells and performed allograft assay with the clonal Non-SMC-GFP cell lines (**Fig. 2D**). Similarly, we found that these subcutaneous tumors also displayed the NCAM^hi^CD44^lo/−^ pattern, ranging from 14±2% to 20±3% (**Fig. 2E and Table S2**). Mixed growth pattern was also observed in culture (**Fig. 2D**). We further isolated the transitioned SMC with NCAM^hi^CD44^lo/−^ pattern and tested their metastasis capability using allograft assay. In contrast to no overt metastasis in the Non-SMC group, multiple distant organ metastases, e.g., lymph node, lung and liver metastases, were detectable in the transitioned SMC group (**Fig. 2F-G and Fig. S2B**). We found that the liver metastases also displayed the NCAM^hi^CD44^lo/−^ pattern (**Fig. 2G**). These data together convincingly proved the transition from Non-SMC to SMC and highlighted the important role of such phenotypic transition in SCLC metastasis.

**Figure 2.**
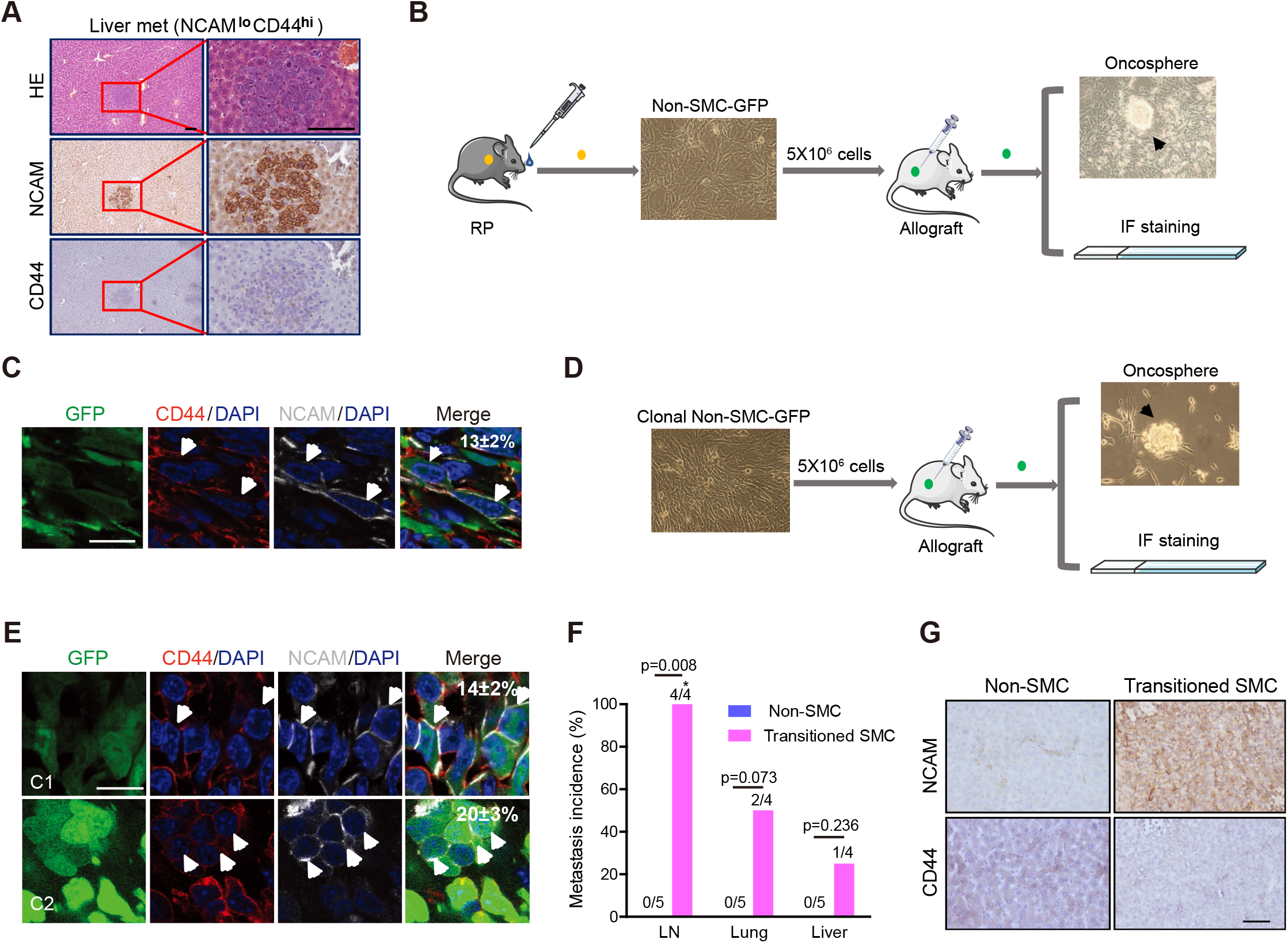
Phenotypic transition from Non-SMC to SMC contributes to SCLC metastasis. **(A)** Representative photos for HE staining, NCAM and CD44 IHC staining in the only one liver metastasis from nude mice subcutaneously transplanted with the NCAM^lo^CD44^hi^ cell lines (Non-SMC) derived from RP mouse model. Scale bars, 100 μm. **(B)** Experimental scheme to test phenotypic transition from Non-SMC to SMC. Primary Non-SMC was derived from RP mouse model and ectopically expressed GFP, and then used for subcutaneous transplantation in nude mice. The subcutaneous tumors were analyzed through onco-sphere formation, and NCAM and CD44 IF staining. Onco-spheres in cell culture were indicated. **(C)** Representative photos of NCAM and CD44 IF staining in subcutaneous tumors from nude mice transplanted with Non-SMC-GFP cells. The NCAM^hi^CD44^lo/−^ subpopulation indicated by white arrow were microscopically counted and the ratio of NCAM^hi^CD44^lo/−^ cells was indicated on the top right corner. Scale bar, 25 μm. Data were shown as mean ± S.E.M. **(D)** Experimental scheme to test the potential phenotypic transition using single cell-derived clonal Non-SMC-GFP. The subcutaneous tumors were then analyzed through onco-sphere formation and NCAM and CD44 IF staining. Onco-spheres in cell culture were indicated. **(E)** Representative photos of NCAM and CD44 IF staining in clonal Non-SMC-GFP subcutaneous tumors. C1: clone #1; C2: clone #2. The NCAM^hi^CD44^lo/−^ subpopulation indicated by white arrow were microscopically counted and the ratio of NCAM^hi^CD44^lo/−^ cells was indicated on the top right corner. Scale bar, 25 μm. Data were shown as mean ± S.E.M. **(F)** Statistical analyses of the incidence of lymph node (LN), lung and liver metastases in nude mice subcutaneously transplanted with transitioned SMC or Non-SMC, which were derived from the clonal Non-SMC-GFP subcutaneous tumors. n=4 mice for transitioned SMC group and n=5 mice for paired Non-SMC group. P values were calculated by Pearson chi-square test. **(G)** Representative photos of NCAM and CD44 IHC staining of mouse livers in **F**. The livers from paired Non-SMC showed no metastasis. Scale bar, 100 μm.

### Brg1 knockout inhibits SMC phenotypic transition and SCLC metastasis

To further explore the molecular mechanisms underlying Non-SMC-to-SMC transition, we performed RNA Sequencing and comparatively analyzed the gene expression profiling of SMC and Non-SMC. Small cell neuroendocrine (SCN) signature is recently established as an important index for SCLC metastasis ^38^. Interestingly, we found a significant enrichment of SCN signature-related pathways in SMC whereas non-SCN related pathways (immune-related pathways) were enriched in Non-SMC (**Fig. 3A, Table S3-4**). Real-time PCR data further confirmed the increased expression of SCN signature genes including *Ascl1*, *Insm1*, *Neurod1*, *Chga*, *Sox11* and *Ttf1* in SMC (**Fig. 3B**). These data might partially explain the high metastasis capability of SMC.

**Figure 3.**
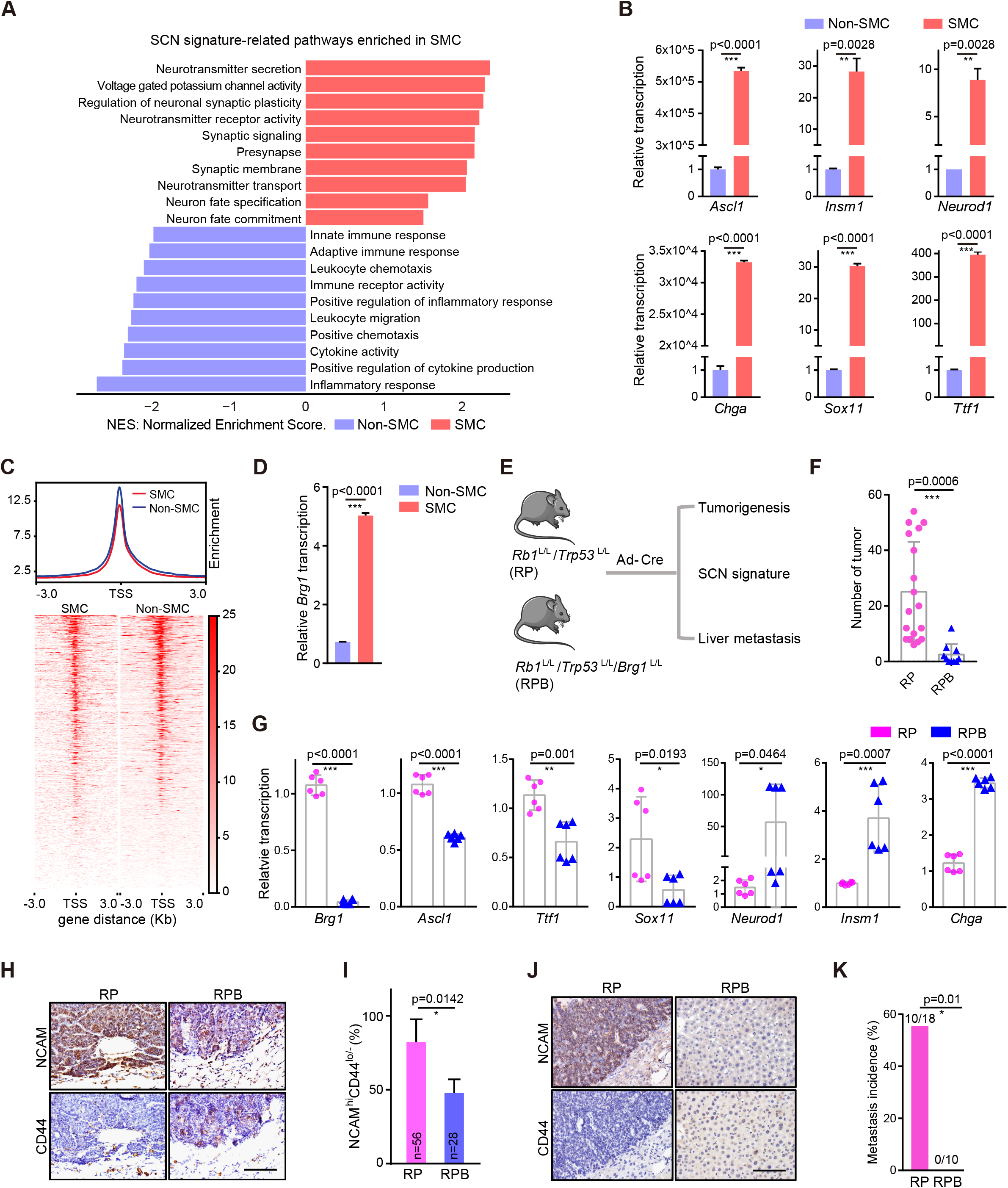
Knockout of *Brg1* in RP mouse model significantly abrogates SMC phenotypic transition and SCLC metastasis. **(A)** The enrichment of small cell neuroendocrine (SCN) signature-related pathways in SMC and the enrichment of immune-related pathways in Non-SMC. NES, Normalized Enrichment Score. **(B)** Real-time PCR detection of SCN signature-related genes including *Ascl1*, *Insm1*, *Neurod1*, *Chga*, *Sox11* and *Ttf1* in SMC vs. Non-SMC. Data were shown as mean ± S.E.M. P values were calculated by unpaired two-tailed *t test*. **(C)** Trend plot (top) and heat map (below) showing ATAC-seq signal over 6kb regions centered at the transcription start sites (TSS) in SMC and Non-SMC. **(D)** Real-time PCR detection of *Brg1* expression in SMC vs. Non-SMC. **(E)** Schematic illustration of the comparative analyses of *Rb1*^L/L^/*Trp53*^L/L^ (RP) and *Rb1*^L/L^/*Trp53*^L/L^/*Brg1*^L/L^ (RPB) mice. **(F)** Statistical analyses of primary tumor numbers in RP and RPB mice at 32 weeks after Ad-Cre treatment. n=18 mice for RP group, n=10 mice for RPB group. Data were shown as mean ± S.E.M. P value was calculated by unpaired two-tailed *t test*. **(G)** Real-time PCR detection of *Brg1* and the SCN signature-related genes in primary tumors from RP and RPB mice. n=2 mice for each group. Data shown as mean ± S.E.M. P values were calculated by unpaired two-tailed *t test*. **(H)** Representative photos of NCAM and CD44 IHC staining in primary tumors from RP and RPB mice at 32 weeks after Ad-Cre treatment. Scale bar, 100 μm. **(I)** Statistical analyses of the percentage of primary tumors with NCAM^hi^CD44^lo/−^ expression pattern in RP and RPB mice. The NCAM^hi^CD44^lo/−^ tumors were defined when the lesions contained more than 50% cells showing NCAM^hi^ and CD44^lo/−^ expression. A total of 56 tumors from 3 RP mice and 28 tumors from 4 RPB mice were analyzed. Data were shown as mean ± S.E.M. P value was calculated by unpaired two-tailed *t test*. **(J)** Representative photos of NCAM and CD44 IHC staining in livers of RP and RPB mice. The livers from RPB mice contained no metastasis. Scale bar, 100 μm. **(K)** Liver metastasis incidence in RP and RPB mice at 32 weeks after Ad-Cre treatment. n=18 mice for RP group, n=10 mice for RPB group. P value was calculated by Pearson chi-square test.

Epigenetic alterations have been implicated in cancer plasticity ^39, 40^. We then performed the assay for transposase-accessible chromatin with next-generation sequencing (ATAC-seq) to determine the global chromatin accessibility of SMC and Non-SMC. Our analyses on transcription start sites revealed an overall reduced signal in the active promoter regions of SMC (**Fig. 3C**). Chromatin remodelers such as SWI/SNF complex is critical for regulating chromatin architecture and accessibility ^18^. We found that multiple members of the SWI/SNF complex, including the central catalytic ATPase *Brg1*, were markedly dysregulated between these two subpopulations (**Fig. S3A**). Real-time PCR quantification further confirmed the significant up-regulation of *Brg1* in SMC (**Fig. 3D**).

To test whether *Brg1* is involved in the phenotypic transition and SCLC metastasis, we generated the *Rb1*^L/L^/*Trp53*^L/L^/*Brg1*^L/L^ (RPB) mouse cohort and performed comparative analyses of tumorigenesis, SCN signature enrichment and metastasis in parallel with RP model (**Fig. 3E**). We found that *Brg1* knockout significantly reduced the tumor number (**Fig. 3F and Fig. S3B-C**). Moreover, several SCN signature-related genes including *Ascl1*, *Ttf1* and *Sox11* were significantly down-regulated in RPB tumors (**Fig. 3G and Fig. S3B**). IHC staining of NCAM and CD44 showed that the percentage of primary tumors with SMC expression pattern was also decreased in the RPB group (**Fig. 3H-I, Table S5**). Notably, no liver metastasis was detected in the RPB group in contrast to about 50% incidence in RP model (**Fig. 3J-K**). These data support that the SWI/SNF complex is important for SMC phenotypic transition and SCLC metastasis.

### Epigenetic silencing of TAZ by SWI/SNF complex in SMC

To identify the downstream mediator of SWI/SNF complex in contribution to SCLC phenotypic transition and metastasis, we first constructed the dysregulated transcriptional factor (TF) network through the integrative analyses of RNA-seq and ATAC-seq data as previously described ^41^. We found that *Ascl1* and *Tead2* were top-ranked TFs with highest number of dysregulated target genes in SMC and Non-SMC respectively (**Fig. 4A-B, Fig. S4-5, Table S6**). ASCL1 is known as the pioneering TF that initializes neuronal reprogramming and also included in the SCN biomarker genes ^42^. TEAD family members are important TFs that function with cofactors YAP/TAZ in cancer malignant progression ^30, 43, 44^. GSEA analysis revealed that the Hippo pathway was significantly enriched in Non-SMC (**Fig. 4C**). Moreover, *Taz/Yap* stood out as top hits among the dysregulated components of Hippo pathway (**Fig. 4D**). Using real-time PCR, we further confirmed the decreased expression of *Taz/Yap* in SMC vs. Non-SMC cells (**Fig. 4E, Fig. S3D**).

**Figure 4.**
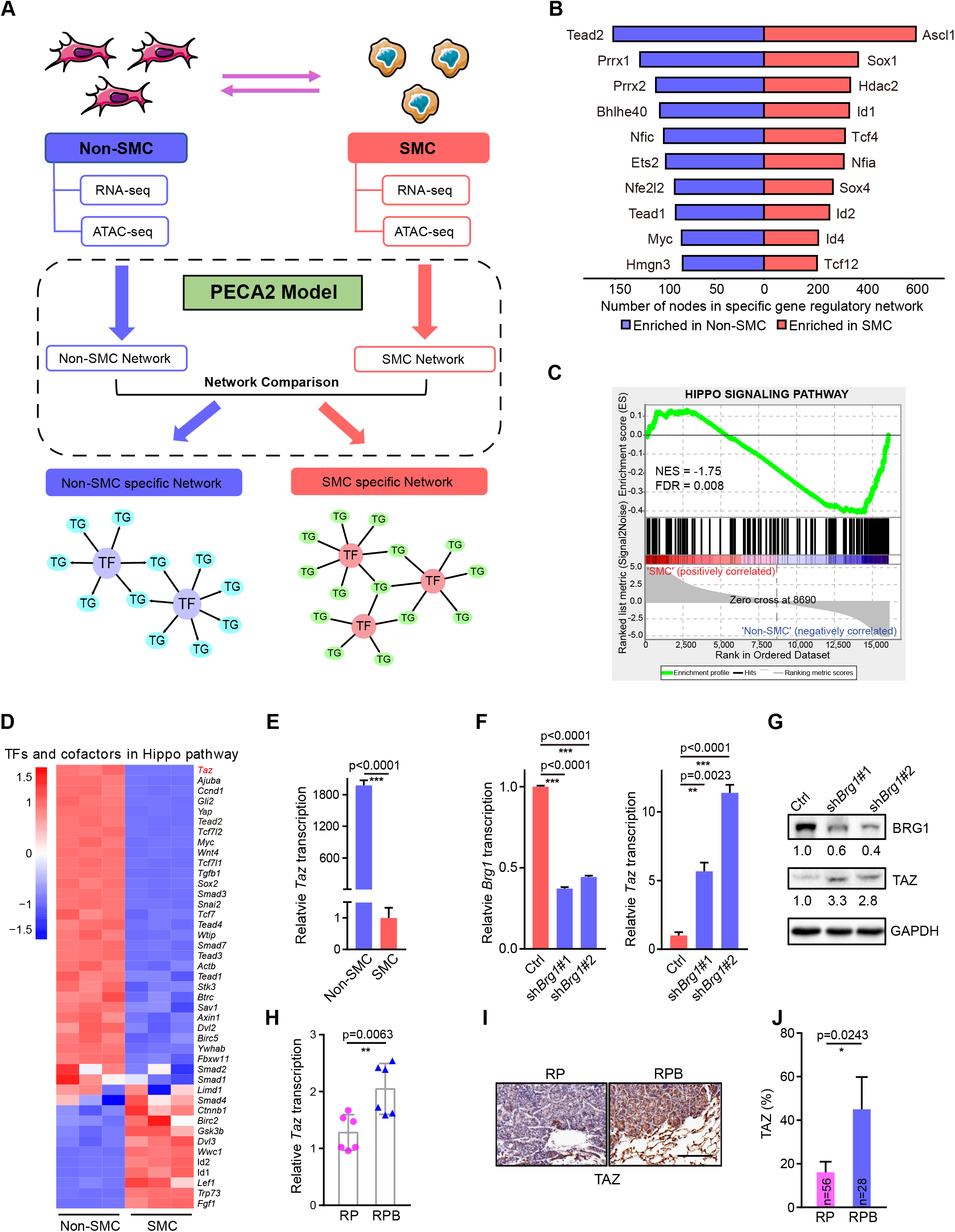
TAZ is epigenetically silenced by the SWI/SNF complex in SMC. **(A)** Schematic illustration of the integrative analyses of RNA-seq and ATAC-seq in SMC and Non-SMC. Specific TF networks in SMC and Non-SMC were constructed according to the PECA2 model (see details in method). **(B)** Enriched transcription factors (TFs) in SMC and Non-SMC through integrative analyses of ATAC-seq and RNA-seq data ranked based on the numbers of dysregulated target genes. **(C)** Gene set enrichment analysis (GSEA) plot of Hippo signaling pathway in SMC vs. Non-SMC. **(D)** Heat map of RNA-seq data showing the relative expression of TFs and cofactors in Hippo pathway in SMC vs. Non-SMC. **(E)** Real-time PCR detection of *Taz* in SMC and Non-SMC. Data were shown as mean ± S.E.M. P value was calculated by unpaired two-tailed *t test*. **(F)** Real-time PCR detection of *Brg1* and *Taz* in SMC with or without *Brg1* knockdown. *Gapdh* served as the internal control. Data were shown as mean ± S.E.M. P values were calculated by unpaired two-tailed *t test*. **(G)** Western blot detection of BRG1 and TAZ levels in SMC with or without *Brg1* knockdown. GAPDH served as the internal control. **(H)** Real-time PCR detection of *Taz* in primary tumors from RP and RPB mice at 32 weeks after Ad-Cre treatment. *Gapdh* served as the internal control. n=2 for each group. Data were shown as mean ± S.E.M. P value was calculated by unpaired two-tailed *t test*. **(I)** Representative photos of TAZ IHC staining in primary tumors from RP and RPB mice at 32 weeks after Ad-Cre treatment. Scale bar, 100 μm. **(J)** Percentage of TAZ positive tumors in RP vs. RPB mice at 32 weeks after Ad-Cre treatment. 56 tumors from 3 RP mice and 28 tumors from 4 RPB mice were analyzed. Data were shown as mean ± S.E.M. P value was calculated by unpaired two-tailed *t test*.

We further asked whether *Taz/Yap* expression was regulated by *Brg1*. We found that *Brg1* knockdown in SMC cells resulted in a significant up-regulation of TAZ expression whereas the expression of YAP was not notably affected (**Fig. 4F-G and Fig. S3E**). Such up-regulation of TAZ was also detectable in RPB tumors in comparison to RP tumors (**Fig. 4H-J, Table S5**). Moreover, we observed a decreased chromatin accessibility at the promoter region of *Taz* in SMC cells (**Fig. S3F**), which might explain the reduced TAZ expression (**Fig. 1D**). In contrast, no substantial change of the chromatin accessibility at the *Yap* promoter region was observed between these two subpopulations (**Fig. S3F**). In supporting of this, TAZ level was obviously down-regulated in primary tumors at extensive stage (**Fig. S1A)**. Moreover, knockdown of *Arid1a* or *Arid2*, another two important components of SWI/SNF complex, obviously up-regulated TAZ expression in SMC cells (**Fig. S3G-H**). However, YAP expression was only slightly up-regulated with *Arid2* knockdown, or even down-regulated after *Arid1a* knockdown in SMC (**Fig. S3G-H**). These results together demonstrate that TAZ is silenced during Non-SMC-to-SMC transition through SWI/SNF complex-mediated epigenetic reprogramming.

### TAZ knockdown promotes Non-SMC-to-SMC transition and accelerates SCLC metastasis

To explore the function of TAZ in phenotype transition and SCLC metastasis, we performed *Taz* knockdown in Non-SMC for allograft assay (**Fig. 5A**). We found that *Taz* knockdown in Non-SMC up-regulated NCAM and down-regulated CD44 *in vitro* (**Fig. 5B and Fig. S6A**). Moreover, *Taz* knockdown also promoted the invasiveness in matrigel, the colony formation in soft agar as well as the anti-anoikis capability in Non-SMC (**Fig. 5C-E**). Immunofluorescence staining of allograft tumors showed that *Taz* knockdown promoted the appearance of NCAM^hi^CD44^lo/−^ pattern, resembling the SMC-derived tumors (**Fig. 5F, Fig. S6B**). Importantly, knockdown of *Taz* in Non-SMC promoted distant organ metastasis (**Fig. 5G**). IHC staining further confirmed that these metastases displayed the SMC expression pattern (**Fig. 5H**). These data together demonstrate that TAZ down-regulation promotes the phenotypic transition from Non-SMC to SMC and SCLC metastasis.

**Figure 5.**
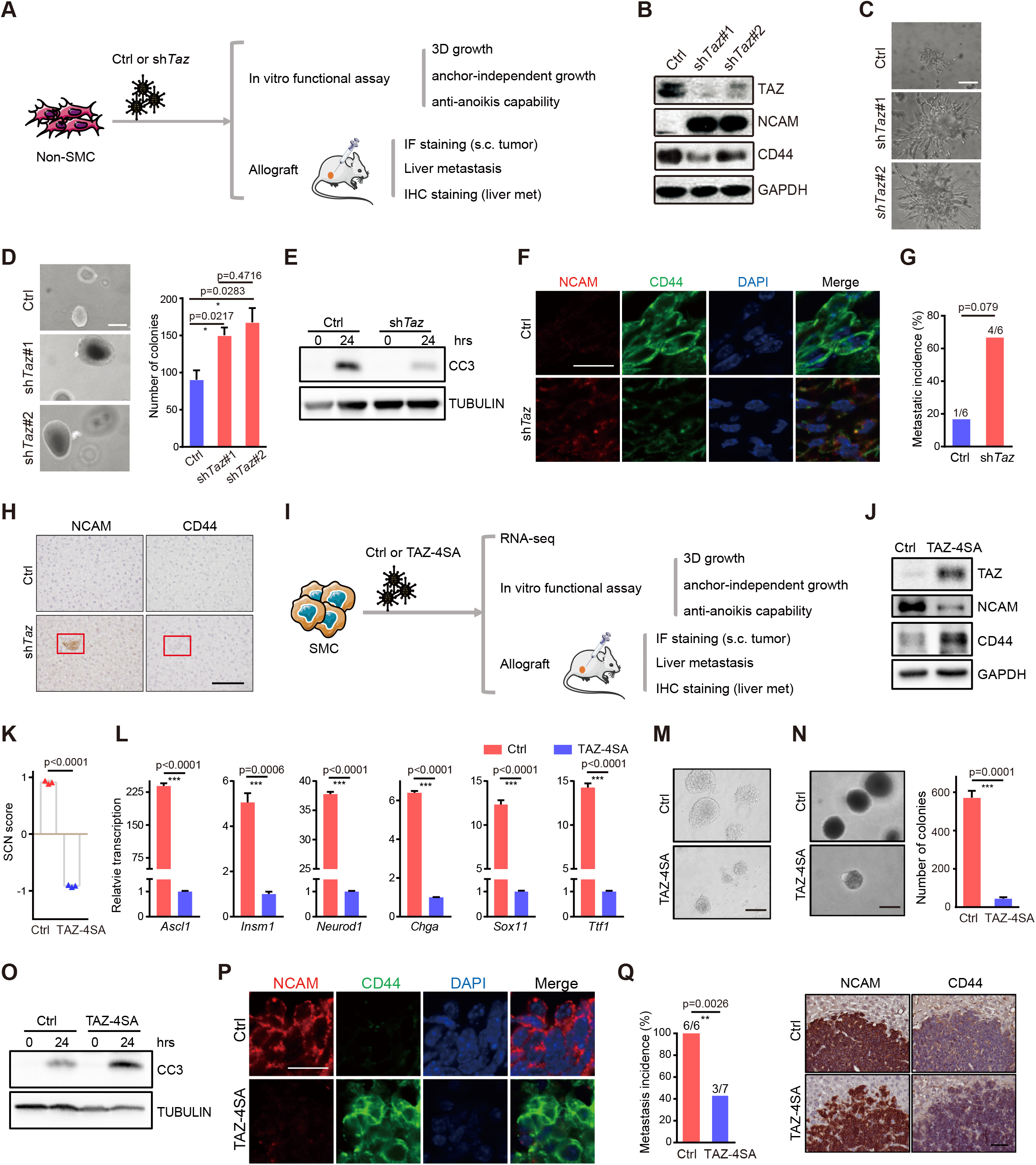
TAZ functions as a critical molecular switch in regulating the phenotypic transition and SCLC metastasis. **(A)** Schematic illustration of the comparative analyses of Non-SMC with or without *Taz* knockdown. **(B)** Western blot detection of TAZ, NCAM and CD44 levels in Non-SMC with or without *Taz* knockdown. GAPDH served as the internal control. **(C)** Matrigel invasiveness of Non-SMC with or without *Taz* knockdown. Scale bars, 100 μm. **(D)** Representative photos (left) and number (right) of the soft-agar colonies of Non-SMC with or without *Taz* knockdown. Scale bars, 100 μm. Data were shown as mean ± S.E.M. P value was calculated by unpaired two-tailed *t test*. **(E)** Western blot detection of cleaved caspase3 (CC3) in anti-anoikis assay of Non-SMC with or without *Taz* knockdown. TUBULIN served as the internal control. **(F)** Representative photos of NCAM and CD44 IF staining in subcutaneous tumors from nude mice transplanted with Non-SMC with or without *Taz* knockdown. Scale bar, 25 μm. **(G-H)** Metastasis incidence (**G**) and representative photos of NCAM and CD44 IHC staining (**H**) in livers from nude mice transplanted by Non-SMC with or without *Taz* knockdown. n=6 for each group. P value was calculated by Pearson chi-square test. Scale bar, 100 μm. **(I)** Schematic illustration of the comparative analyses of SMC with or without ectopic TAZ-4SA expression. **(J)** Western blot detection of TAZ, NCAM and CD44 levels in SMC with or without ectopic TAZ-4SA expression. GAPDH served as the internal control. **(K)** SCN score of SMC with or without ectopic TAZ-4SA expression. Data were shown as mean ± S.E.M. P value was calculated by unpaired two-tailed *t test*. **(L)** Real-time PCR detection of the SCN signature-related genes in SMC with or without ectopic TAZ-4SA expression. *Gapdh* served as the internal control. Data were shown as mean ± S.E.M. P values were calculated by unpaired two-tailed *t test*. **(M)** Matrigel invasiveness of SMC with or without ectopic TAZ-4SA expression. Scale bar, 100 μm. **(N)** Representative photos (left) and statistical analyses (right) of soft-agar colonies of SMC with or without ectopic TAZ-4SA expression. Scale bar, 100 μm. Data were shown as mean ± S.E.M. P value was calculated by unpaired two-tailed *t test*. **(O)** Western blot detection of CC3 in anti-anoikis assay of SMC with or without ectopic TAZ-4SA expression. TUBULIN served as the internal control. **(P)** Representative photos of NCAM and CD44 IF staining in subcutaneous tumors from nude mice transplanted with SMC with or without ectopic TAZ-4SA expression. Scale bar, 25 μm. **(Q)** Metastasis incidence (left) and representative photos of NCAM and CD44 IHC staining of liver metastasis (right) in nude mice transplanted with SMC with or without ectopic TAZ-4SA expression. n=6 mice for Ctrl group, n=7 mice for TAZ-4SA group. Scale bar, 100 μm. P value was calculated by Pearson chi-square test.

### Ectopic TAZ expression reversely promotes the transition from SMC to Non-SMC and alleviates SCLC metastasis

To test if the phenotypic transition from Non-SMC to SMC is reversible, we ectopically expressed a constitutive activated TAZ mutant (TAZ-4SA) ^30^ in SMC (**Fig. 5I**). We found that ectopic TAZ-4SA expression in SMC dramatically down-regulated NCAM and up-regulated CD44 expression *in vitro*, indicative of the potential reversible transition from SMC to Non-SMC (**Fig. 5J, Table S7-8**). Moreover, the SCN score and related genes expression also decreased after ectopic TAZ4SA expression (**Fig. 5K-L, Table S7-8**). Functional assays showed that TAZ-4SA expression markedly suppressed the matrigel invasiveness, colony formation in soft agar, and anti-anoikis capability of SMC (**Fig. 5M-O**). Immunofluorescence staining also showed that ectopic TAZ-4SA expression promoted the Non-SMC expression pattern in comparison to SMC-derived subcutaneous tumors (**Fig. 5P, Fig. S6C**).

More importantly, ectopic TAZ-4SA expression significantly suppressed the liver metastasis of SMC (**Fig. 5Q**). These findings support that ectopic *Taz* expression promotes the reverse transition from SMC to Non-SMC and alleviates SCLC metastasis.

### Low TAZ level associates with SCN signature enrichment and predicts poor prognosis of SCLC patients

To evaluate whether our findings are clinically relevant, we downloaded public RNA sequencing dataset of 112 human SCLC ^2, 45^ and analyzed the correlation between *Taz* and SCN signature, and single-cell sequencing data of liver metastasis ^15^ to detect whether SMC exist in metastatic lesion, and collected 101 Chinese surgical specimens for prognosis analyses (**Fig. 6A**). Bioinformatic analyses showed that human SCLC with low TAZ expression (TAZ^lo^) displayed a significantly higher SCN score (**Fig. 6B, Table S9**), indicative of strong metastasis capability. The SCN signature-related pathways including Positive regulation of neurotransmitter transport, Neurotransmitter secretion, and Synaptic vesicle membrane, were significantly enriched in TAZ^lo^ SCLC (**Fig. S7A**). Consistently, most SCN signature-related genes, including ASCL1, INSM1, and CHGA, were significantly increased in TAZ^lo^ SCLC samples (**Fig. 6C**). Moreover, NCAM was increased, and CD44 was decreased in TAZ^lo^ SCLC specimens (**Fig. 6C**), indicative of the SMC pattern of these samples. And TEAD2 also decreased in these TAZ^lo^ samples (**Fig. 6C**). Besides, we observed a negative correlation between the SCN signature-related genes and TAZ, and positive correlation between CD44, TEAD2 and TAZ (**Fig. S7B, Table S9)**.

**Figure 6.**
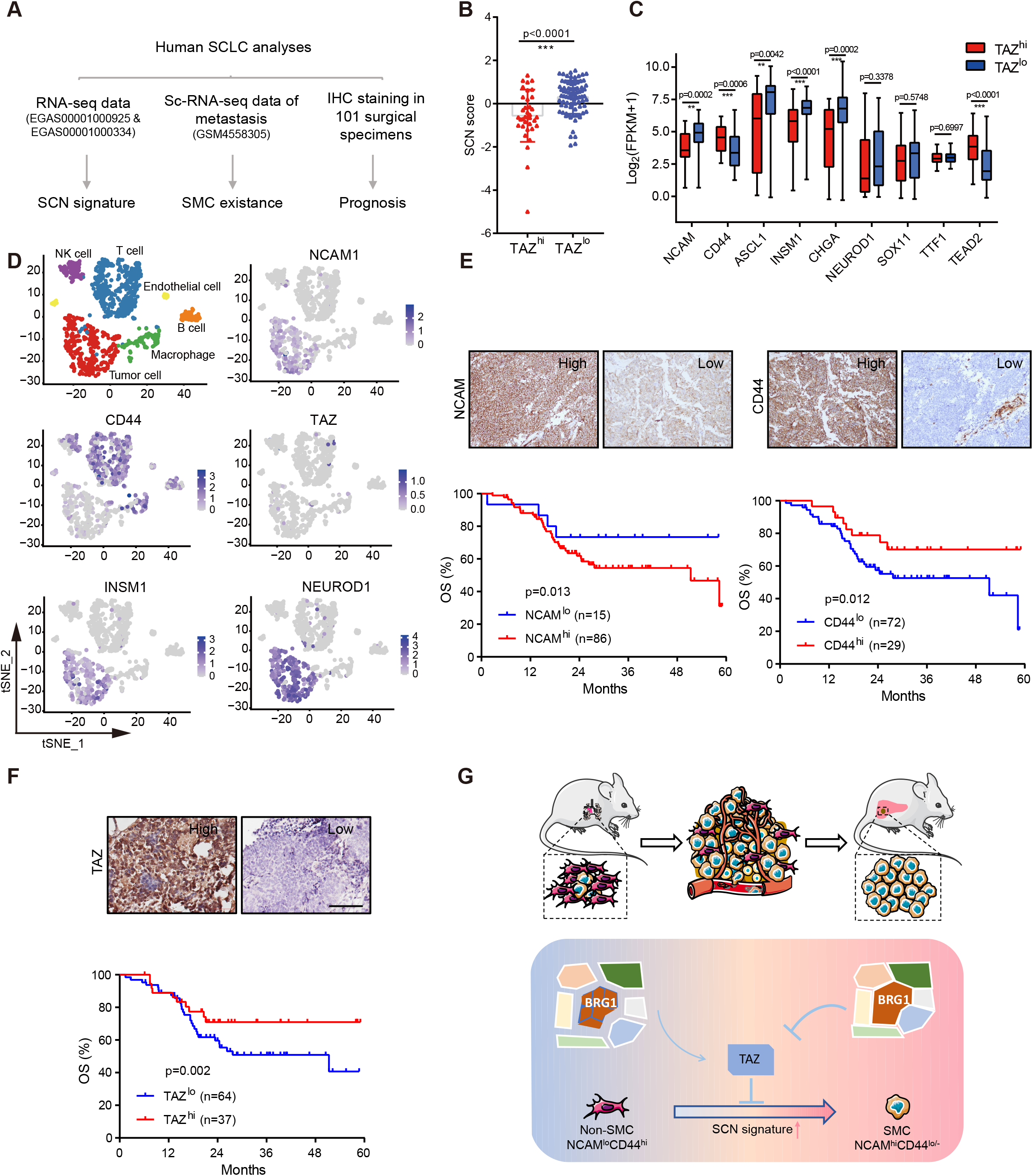
Low TAZ level is correlated with SCN signature enrichment and predicts poor prognosis of SCLC patients. **(A)** Schematic illustration of the analyses of human SCLC specimens. **(B)** SCN score of human SCLC specimens with high or low TAZ mRNA level. The RNA-seq data were down-loaded from public database (GSE69091 and EGAS00001000334). Data were shown as mean ± S.E.M. P value was calculated by unpaired two-tailed *t* test. **(C)** Correlation between individual SCN signature-related genes, CD44 or TEAD2 expression with high or low TAZ level in human SCLC (GSE69091 and EGAS00001000334). Data were shown as mean ± S.E.M. P values were calculated by unpaired two-tailed *t* test. **(D)** Clustering and the NCAM, CD44, TAZ, NEUROD1 and INSM1 expression of the single cell sequencing data (GSM4558305) of liver biopsy from SCLC patient. **(E-F)** Representative photos of NCAM, CD44 (**E**) or TAZ (**F**) IHC staining in Chinese SCLC specimens (up) and survival curves of high or low expression of NCAM, CD44 or TAZ with overall survival (OS) (below). Scale bar, 100 μm. P values were calculated by Kaplan-Meier analysis with log-rank test. **(G)** Working model illustrating the essential role of SWI/SNF complex-mediated TAZ expression in controlling the phenotypic transition from Non-SMC to SMC and SCLC metastasis. TAZ, which is epigenetically silenced by the SWI/SNF complex, functions as a critical molecular switch during the phenotypic transition from Non-SMC to SMC and SCLC metastasis. Disruption of SWI/SNF complex through BRG1 knockout promotes TAZ up-regulation and thus inhibits the phenotypic transition and cancer metastasis.

We further took advantage of the Ireland et al. single-cell RNA sequencing data derived from SCLC liver metastasis ^15^. Interestingly, we found that most SCLC metastatic cells showed high expression of NCAM with concurrent low expression of CD44, resembling the SMC pattern (**Fig. 6D**). Moreover, these cells showed high expression of the SCN signature markers INSM1 and NEUROD1, similar to the SMC in RP model (**Fig. 6D**). Importantly, low or no TAZ expression was detected in these metastasis cells (**Fig. 6D**), confirming the silence of TAZ in metastasis.

Lastly, we collected a patient cohort containing 101 Chinese SCLC surgical specimens for immunostaining analyses of NCAM, CD44 and TAZ. Most of these patients were at limited stage without distant metastases. We found that high NCAM or low CD44 levels were significantly associated with worse patient overall survival (OS) (**Fig. 6E**, **Table S10**). Moreover, TAZ^lo^ patients also showed a worse overall survival (**Fig. 6F**). These data together provide strong clinical evidence in support of our findings of SMC in RP model.

## Discussion

SCLC is the most lethal form of lung cancer, characterized with highly metastatic capacity. A growing body of evidence based on mouse models has demonstrated that SCLC is highly heterogeneous with distinct subpopulations playing different roles during malignant progression and metastasis ^5–15^. In this study, we identify the NCAM^hi^CD44^lo/−^ cells in RP model as the SCLC metastasizing cells. We further reveal that the SMCs are progressively transitioned from Non-SMCs during SCLC malignant progression and metastasis. Our data further show that the SWI/SNF complex-mediated epigenetic down-regulation of TAZ is essential for driving such phenotype transition. Moreover, TAZ activation is sufficient to drive the reverse transition from SMC to Non-SMC and thus alleviates SCLC metastasis. With the support from clinical specimen analyses, our data demonstrate that the NCAM^hi^CD44^lo/−^ cells are mainly responsible for SCLC metastasis and the SWI/SNF-TAZ axis importantly orchestrates SCLC plasticity and metastasis (**Fig. 7G**).

To assess the SCLC heterogeneity in RP model, we use both NE marker NCAM and mesenchymal marker CD44 to do the immunostaining and FACS analyses, and identify the NCAM^hi^CD44^lo/−^ cells as the SCLC metastasizing cells. Previous study shows that mouse SCLC cells contain both NE and non-NE subpopulations ^9^. However, neither subpopulation alone can metastasize and a synergetic cooperation is necessary for distant organ metastasis ^9^. In contrast, our data show that the NCAM^hi^CD44^lo/−^ cells harbor strong metastasis capability in allograft assay, and the tumors metastasize into multiple distant organs including lymph node, lung and liver. Since the SMC defined here also express the classical NE biomarkers, we reason that the NCAM^hi^CD44^lo/−^ cells might belong to the NE subpopulation, but with higher metastasis potential. In another words, the NCAM^hi^CD44^lo/−^ cells might represent as the highly metastatic subpopulation of the NE subtype. Future efforts looking into the heterogeneity of NE subtype will hopefully uncover more subpopulations in link to SCLC malignant progression and metastasis.

We also find that phenotypic transition from Non-SMC to SMC contributes to SCLC metastasis, which closely links cancer plasticity and malignant progression. Indeed, recent data also show, during SCLC drug resistance acquisition, that Notch signaling promotes the transition from NE to non-NE subtype and thus provides a niche for resisting to drug treatment ^10^. Similar transition from the NE to non-NE subtypes have also been found in another recent study ^15^. Metastasis and drug resistance are two major hurdles in clinical SCLC management. Understanding of molecular mechanisms involved in the phenotypic transition in these two important events will hopefully provide a solid base for the development of novel therapeutic strategy to treat SCLC in the clinic.

Through integrative analyses of gene expression profiling and chromatin accessibility, we find that the SWI/SNF complexes play an important role during Non-SMC to SMC transition. Knockout of its ATPase BRG1 inhibits such phenotypic transition and cancer metastasis, indicating an oncogenic function of SWI/SNF complex as well as BRG1 in SCLC. Previous study shows that BRG1 is important for the activation of neuroendocrine transcription ^25^. Consistently, we observe the suppression of neuronal gene expression by BRG1 knockdown and the enrichment of SCN signature in SMC subpopulation.

We further find that TAZ is an important downstream mediator of SWI/SNF complex during SCLC phenotypic transition. Although both YAP and TAZ are significantly up-regulated in Non-SMC, only TAZ is significantly up-regulated when *Brg1* is knocked down in SMC. Similar findings are also observed when *Arid1a* or *Arid2* is knocked down. Consistently, *Brg1* knockout in RP mouse up-regulates TAZ and significantly inhibits the SMC appearance and SCLC metastasis. Moreover, we find that low TAZ expression is associated with the SCN signature enrichment. In agreement with these observations, previous studies have shown that high YAP/TAZ expression correlates with decreased NE markers ^46^, and YAP loss defines NE differentiation ^47^. Meanwhile, NE lineage markers are dominant in the SCN signature, which significantly associates with SCLC malignant progression and metastasis ^38,48–50^. Of course, considering the redundant function between YAP and TAZ, it remains possible that YAP might also regulate SCLC phenotypic transition independent of SWI/SNF complex. Future efforts will be necessary to clarify detailed regulatory mechanisms underlying YAP expression during SCLC phenotypic transition and metastasis.

Our findings from loss-of-function and gain-of-function experiments support a tumor suppressive role of TAZ in SCLC. YAP/TAZ are well established as oncogenic drivers. Nonetheless, accumulating evidence has recently revealed a tumor suppressive function of YAP/TAZ in multiple cancer types ^51^. For instance, YAP resitrics Wnt signals during intestinal regeneration which results in rapid loss of intestinal crypts, and YAP loss promotes hyperplasia and microadenomas development ^52^. In hematological cancer, low YAP level prevents nuclear ABL1-induced apoptosis and rescued YAP expression triggers cell death ^53^. Another study shows that the growth inhibitory effect caused by LATS1/2 deletion is due to uncontrolled activation of YAP in colon cancer ^54^. A latest report demonstrates that LATS1/2 promotes breast cancer cell growth through inhibition of YAP/TAZ ^34^. Our findings of the tumor-suppressive function of TAZ are also supported by clinical specimen analyses. Single cell RNA sequencing data support that the cancer cells from SCLC liver metastasis mainly display the SMC expression pattern and these metastatic cells show low or no expression of TAZ. Moreover, low TAZ level is significantly associated with poor patient survival. These data together support that TAZ works as a tumor suppressor in controlling SCLC plasticity and metastasis.

## Materials and Methods

### RP and RPB mouse cohort generation, maintenance and analyses

Mice were housed in a specific pathogen-free environment at the Shanghai Institute of Biochemistry and Cell Biology, and treated in accordance with protocols conformed to the ARRIVE guidelines and approved by the Institutional Animal Care and Use Committee of the Shanghai Institutes for Biological Sciences, Chinese Academy of Sciences (approval number: IBCB0011). Conditional knockout mice including *Trp53*^L/L^, *Rb1*^L/L 3^ and *Brg1*^L/L 55^ alleles were generously provided by Drs. Tyler Jacks, Ronald A. DePinho and Pierre Chambon. Mice were crossed to obtain *Rb1*^L/L^/*Trp53*^L/L^ (RP) and *Rb1*^L/L^/*Trp53*^L/L^/*Brg1*^L/L^ (RPB) cohorts. All experimental mice were maintained on a mixed genetic background as previously described ^56^. Mice at 6-8 weeks old were treated with Adenovirus-CMV-Cre recombinase (Ad-Cre, 2×10^6^ p.f.u.) by intratracheal intubation ^57^ to allow for Cre-lox mediated recombination of floxed alleles. Mouse tumors were used for immunostaining, FACS analyses, genomic DNA extraction and genotyping as previously described ^3, 55^. The primers sequences are shown in supplementary data.

### Statistical analysis

Statistical analyses were carried out using SPSS 16.0 or GraphPad Prism 5/7 software (San Diego, CA). The significance of differences was determined using two-tailed Student’s *t* test or chi-square test. Kaplan-Meier analysis with log-rank test was used to assess patients’ survival between subgroups. P value < 0.05 was considered to be statistically significant.

## Data availability

Sequence data have been deposited in GEO with the primary accession code GSE158091 (ATAC-seq of SMC and Non-SMC), GSE158290 (RNA-seq of SMC and Non-SMC) and GSE158293 (RNA-seq of SMC-Ctrl and SMC-TAZ-4SA).

## Acknowledgements

We thank Drs. Tyler Jacks, Ronald A. DePinho for RP mouse model and Dr. Pierre Chambon for *Brg1^L/L^* mouse. We are grateful to Drs. Fuming Li, Xiangkun Han for technical assistance and Drs. Carla F Kim, Dangsheng Li, Cheng Li, Yujiang Geno Shi, Rui Fang and Nella Dost for constructive comments.

## Author Contributions

H.J. and Y.J. conceived the idea and designed the experiments. Y.J., Q.Z., Y.F., T.X., and H.H. performed all experiments and analyzed the data. W.Z., Y.W., J.C., Y.C. and L.C. performed the bioinformatics analyses. P.Z., L.J. and Y.H. provided human SCLC specimens. C.G., K.K.W., R.K.T., H.Z., X.Z., D.G., J.Q., F.L., and X.Y.L. provided technical assistance and helpful comments. H.J., Y.J. and L.H. wrote the manuscript.

