## Supplemental data for "Identification of TAZ as the essential molecular switch in orchestrating SCLC phenotypic transition and metastasis"

^13^Department of Translational Genomics, Center of Integrated Oncology Cologne-Bonn, Medical Faculty, University of Cologne, 50931 Cologne, Germany. Department of Pathology, University Hospital Cologne, 50937 Cologne, Germany.

^14^School of Life Science, Hangzhou Institute for Advanced Study, University of Chinese Academy of Sciences, Hangzhou 310024, China

^15^Leading author.

^16^These authors contributed equally to this work.

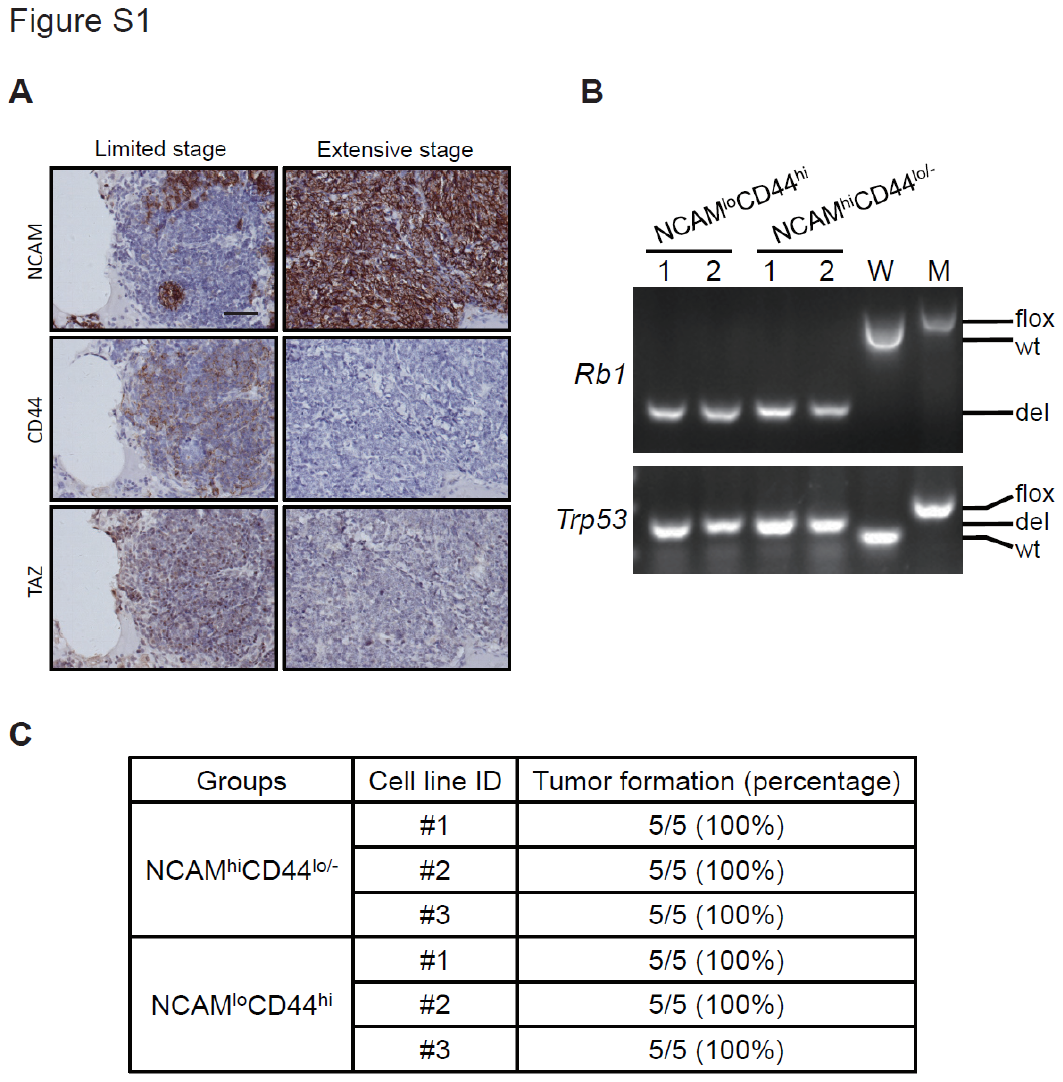

**Supplementary Figure 1. Identification of the NCAM^hi^CD44^lo/-^ subpopulation as SCLC metastasizing cells in RP mouse model.**

**(A)** Representative photos of NCAM, CD44 and TAZ IHC staining of primary tumors at limited stage (no overt metastasis) and extensive stage (overt metastasis) in RP model. Scale bar, 100 μm. **(B)** Genotyping of *Rb1* and *Trp53* in NCAM^lo^CD44^hi^ and NCAM^hi^CD44^lo/-^ cells. wt, wild-type allele. flox, conditional knockout allele. del, knockout allele. W, negative control for wild-type. M, positive control for homozygous conditional knockout. (**C**) Tumor formation incidence in allograft assay of 5x10^6^ NCAM^lo^CD44^hi^ and NCAM^hi^CD44^lo/-^ cells in nude mice.

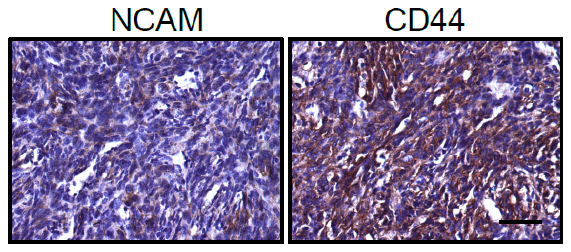

**Supplementary Figure 2.** Representative photos of NCAM and CD44 IHC staining in the subcutaneous tumors from Non-SMC allograft assay.

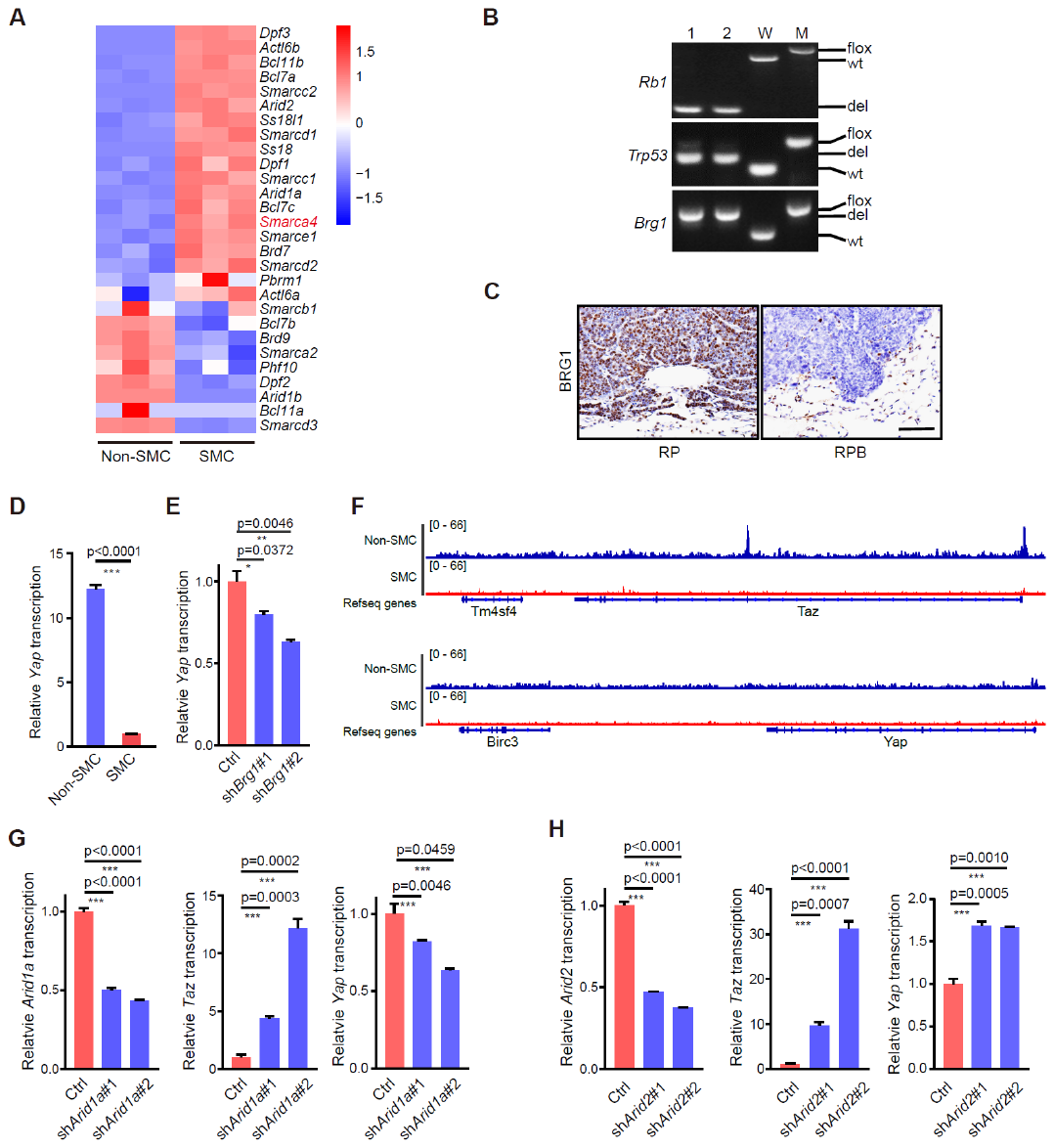

**Supplementary Figure 3. TAZ is epigenetically silenced by SWI/SNF complex in SMC.**

**(A)** Heatmap of RNA-seq data showing differentially expressed components of SWI/SNF complex in SMC vs. Non-SMC. *Smarca4* (*Brg1*) was indicated in red. **(B)** Genotyping of *Rb1*, *Trp53* and *Brg1* alleles in primary lung tumors from RPB mouse model. wt, wild-type allele. flox, conditional knockout allele. del, knockout allele. W, negative control for wild-type. M, positive control for homozygous conditional knockout. **(C)** Representative photos of BRG1 IHC staining in primary lung tumors from RP and RPB models. Scale bar, 100 μm. **(D)** Real-time PCR detection of *Yap* in SMC vs. Non-SMC. Data were shown as mean ± S.E.M. P value was calculated by unpaired two-tailed *t* test. **(E)** Real-time PCR detection of *Yap* in SMC with or without *Brg1* knockdown. Data were shown as mean ± S.E.M. P value was calculated by unpaired two-tailed *t* test. **(F)** Genome browser view of ATAC-seq signal showing peak location of *Taz* and *Yap* in SMC (red) and Non-SMC (blue). **(G)** Real-time PCR detection of *Arid1a, Taz* and *Yap* in SMC with or without *Arid1a* knockdown. Data were shown as mean ± S.E.M. P values were calculated by unpaired two-tailed *t* test. **(H)** Real-time PCR detection of *Arid2*, *Taz* and *Yap* levels in SMC with or without *Arid2* knockdown. Data were shown as mean ± S.E.M. P values were calculated by unpaired two-tailed *t* test.

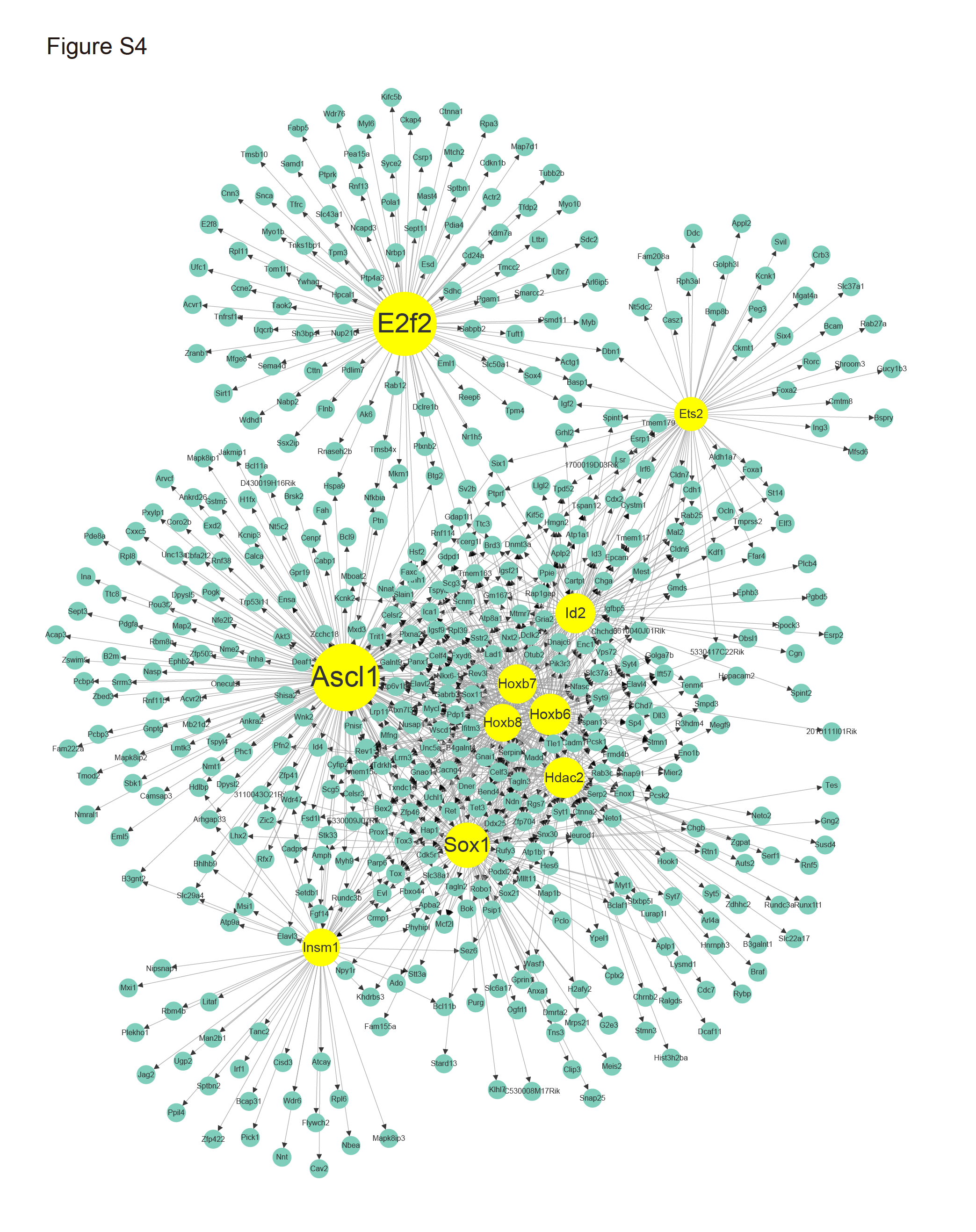

**Supplementary Figure 4. Enriched transcription factor network in SMC through integrative analyses of ATAC-seq and RNA-seq data.** Size of TF node represented the number of its downstream target genes in constructed network. Top 10 TFs were highlighted in yellow which held the largest number of dysregulated genes.

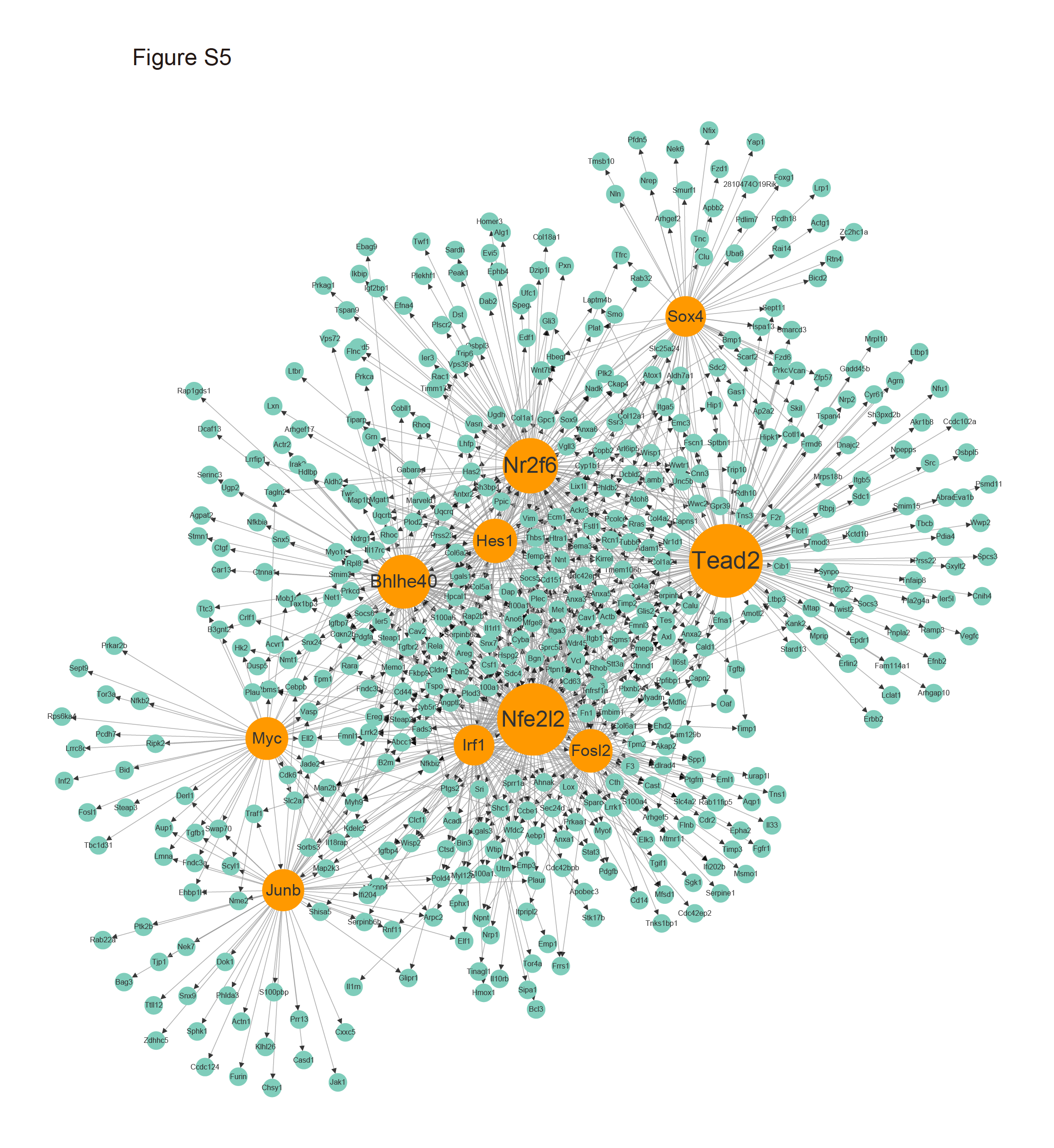

**Supplementary Figure 5.** **Enriched transcription factor network in Non-SMC through integrative analyses of ATAC-seq and RNA-seq data.** Size of TF node represented the number of its downstream target genes in constructed network. Top 10 TFs were highlighted in orange which held the largest number of dysregulated genes.

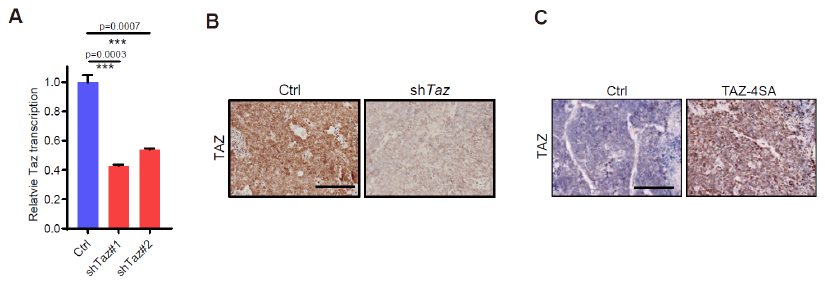

**Supplementary Figure 6.** **Validation of TAZ expression in Non-SMC-sh*Taz* and and SMC-TAZ-4SA in subcutaneous tumors from allograft assay.**

**(A)** Real-time PCR quantification of *Taz* expression in Non-SMC-sh*Taz* cells. P values were calculated by two-tail paired *t* test. **(B)** Representative photos of TAZ immunostaining in Non-SMC and Non-SMC-sh*Taz* subcutaneous tumors in allograft assay. Scale bar, 100 μm. **(C)** Representative photos of TAZ immunostaining in SMC and SMC-*TAZ-4SA* subcutaneous tumors in allograft assay. Scale bar, 100 μm.

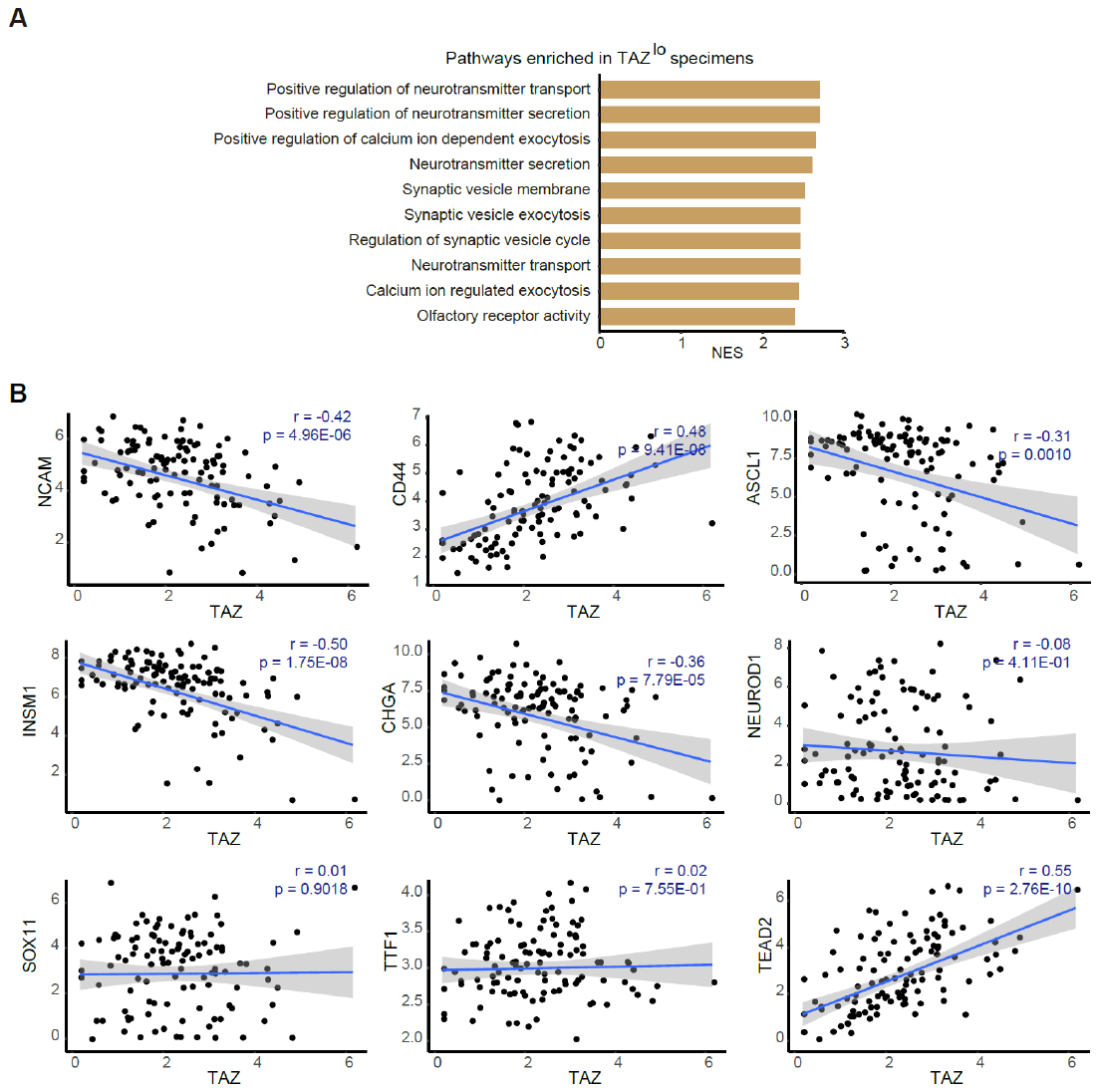

**Supplementary Figure 7. L****ow TAZ level is correlated with high score of SCN signature.**

**(A)** Enrichment of small cell neuroendocrine (SCN) signature-related pathways in human SCLC specimens with low TAZ expression (TAZ^lo^). NES, Normalized Enrichment Score. **(B)** The Pearson correlation coefficient *r* between TAZ and NCAM, CD44, SCN signature-related genes and TEAD2 expression (log_2_(FPKM+1)). The RNA-seq data used in **A** and **B** were downloaded from public database with accession of EGAS00001000925 and EGAS00001000334.

**Supplemental Experimental Procedures**

**RP and RPB mouse model studies**

RP and RPB tumors were stained with NCAM (CST, #99746), CD44 (Santa Cruz, #SC-18849), TAZ (CST, #4883S) and BRG1 (Abcam, #ab110641) antibodies, and blindly scored as high or low expression dependent on the membrane staining densities as described previously ^1^. Antigen retrieval was performed by microwave in sodium citrate (pH 6.0) or EDTA (pH 8.0) buffer. Primary tumors are considered as the NCAM^hi^CD44^lo/-^ expression pattern when more than 50% of cancer cells expressed high NCAM and low/no CD44 as previously described ^2^.

**Histological analysis**

Tumor tissues were dissected and fixed in 4% formaldehyde overnight, and then dehydrated in ethanol, embedded in paraffin, and sectioned (5 μm) followed by staining with hematoxylin and eosin.

**SMC, Non-SMC and single-cell clone establishment**

Primary RP lung tumors or subcutaneous tumors from allograft assay were dissected, dissociated, stained with anti-NCAM-FITC (R&D, FAB7820G) and anti-CD44-APC (eBioscience, E07145-1631) and subjected to FACS analyses using BD Aria II SORP following manufacture’s protocol. The NCAM^hi^CD44^lo/-^ and NCAM^lo^CD44^hi^ subpopulations were isolated and used for the following experiments. These isolated cells were also used for genotyping. Non-SMCs were transfected with GFP (Non-SMC-GFP) and single cells were picked up for the establishment of clonal Non-SMC-GFP stable cell lines. All primary cells or stable cell lines were cultured in RPMI1640 plus 10% FBS and 1% penicillin/streptomycin and free of mycoplasma contamination.

**Allograft transplantation in nude mice**

A total of 5×10^6^ cells were subcutaneously transplanted into nude mice and distant organ metastases were analyzed 10 weeks later. Tumor volumes were monitored twice a week. Subcutaneous tumors and/or distant organ metastases were stained with NCAM (CST, #99746), CD44 (Santa Cruz, #sc-18849) and TAZ (CST, #4883S) antibodies. We also performed immunofluorescence staining in subcutaneous tumors using anti-NCAM (CST, #99746) and anti-CD44 (Santa Cruz, #sc-18849). In brief, tissues were fixed in 4% paraformaldehyde (PFA) overnight, washed in 1×PBS, dehydrated in 25% and 35% sucrose solution serially, then embedded in Tissue Freezing Medium (Leica) and sectioned (5 μm) before staining. Alexa Fluor 533 goat anti-rat or Alexa Fluor 647 goat anti-rabbit was used as secondary antibodies for immunofluorescence staining. Photos were taken using confocal microscopy (Leica TCS SP5). Ten fields per tumor were used for calculating the percentage of cells with NCAM^hi^CD44^lo/-^ expression pattern.

**RNA-Seq library preparation and data processing**

Total RNAs from SMC or Non-SMC, and SMC-Ctrl or SMC-TAZ-4SA were extracted using the TRIzol reagent (Life Technologies) according to the manufacturer’s instruction. All sequencing reactions were performed on an Illumina NovaSeq6000 platform (Berry Genomics Corporation, China). The sequence mode is 150 bp paired-end. We mapped RNA-seq reads to the mouse reference genome mm10 with STAR v2.6.1b. The differential expression genes were detected by DESeq2, and GSEA was performed following the developer’s protocol (http://www.broad.mit.edu/gsea/).

**ATAC-Seq analysis**

Cells were collected and lysed with lysis buffer (10 mM Tris·Cl (pH 7.4), 10 mM NaCl, 3 mM MgCl_2_，0.1% (v/v) NP40), and then fragmented by TruePrepTM DNA Library Prep Kit V2 (Vazyme TD501). All sequencing reactions were performed on an Illumina HiSeq X Ten platform and 150 bp paired-end reads were generated (Berry company). We mapped ATAC-seq reads to the mouse reference genome mm10 with Bowtie2 v2.3.4, and converted output SAM files from Bowtie2 to BAM files using samtools v1.4 for BAM files sorting and indexing. We removed duplicate reads by sambamba v0.6.6 and called peaks using MACS2 v2.1.1. Enrichment analysis of motif and transcription factor were performed with HOMER v4.9. We performed peak annotation with ChIPseeker and used Deeptools for data visualization.

**Transcription factor network analyses**

Transcription factor (TF) network analyses with integrative information from RNA-Seq and ATAC seq were performed as described previously ^3^. We used paired expression and chromatin accessibility (PECA) gene regulatory network to analyze the TF network in SMC and Non-SMC. By taking paired expression (RNA-seq) and chromatin accessibility (ATAC-seq) data as inputs, we employed PECA2 model to generate gene regulatory network in each sample. The input data of PECA2 was paired and processed expression data of RNA-seq and bam file of ATAC-seq. The output contained TF-RE-TG gene regulatory network and detailed statistics information about RE and TF-TG pair. We used PECA_compare_dif.sh to do comparison between generated networks and obtain the specific regulatory network, and counted the nodes of TF in each network and ranked them accordingly.

**SCN score analysis**

SCN score was calculated through projecting RNA-seq data to the varimax-rotated PCA results as previously described ^4^. The Zscored_SCN_score was used for the comparative analyses. Beside the analyses of SMC and Non-SMC, we also used human SCLC RNA-Seq datasets for SCN score analyses, which contained 112 samples including 81 from EGAS00001000925 (European Genome-phenome Archive under the accession code) and 31 from EGAS00001000334 (European Bioinformatics Institute, EBI) ^5^^,^^6^. We performed combat function from package ‘sva’ in R to remove batch effect, and divided samples into TAZ^hi^ or TAZ^lo^ group with the expression cutoff of 3. Using MSigDB C5 gene sets, we performed GSEA on the expression data of the groups, and used pearson correlation coefficient to indicate the gene-gene correlation.

**Single-cell transcriptome analyses of human SCLC liver metastasis**

We used Seurat, an R package (v4.0.1), for the analyses of single cell RNA-seq dataset from human SCLC liver metastasis (GEO: GSM4558305) ^7^. We discarded those cells with less than 300 expressing genes or with more than 7,000 expressing genes (potentially cell doublets), and filtered the genes expressed in less than three cells. In total, we obtained 1,249 cells with mitochondrial gene percentage at less than 10%. The UMI counts were transformed and normalized using the ‘‘NormalizeData()’’ function with default parameters. Principle component analysis (PCA) was performed using the highly variable genes that were identified by function ‘‘FindVariableGenes()’’. The first twenty principal components were applied to perform t-SNE. Cell populations were identified using the ‘‘FindClusters()’’ function with resolution set to 0.3, and clusters were defined on the expression of cell-type-specific gene signatures (for tumor markers, endothelial cells, T cells, B cells, NK cells and Macrophage).

**Analyses of Chinese SCLC specimens**

A total of 101 human SCLC surgical specimens were collected from 2007 to 2012 with the approval by the institutional review committees of Shanghai Pulmonary Hospital (Tongji University), Shanghai Chest Hospital (Jiaotong University), and Zhongshan Hospital (Fudan University). Patients gave written informed consents. All the cases were re-reviewed and confirmed to be SCLC by the pathologists. Most patients (98 out of 101) were diagnosed with limited-stage disease without distant metastases. The specimens were serially sectioned and stained with NCAM (DAKO, #M7304), CD44 (Santa Cruz, #sc-18849) and TAZ (CST, #4883S) antibodies. The IHC staining was blindly scored as high or low according to staining density and subjected to clinical relevance analyses as described previously ^1^.

**Ectopic gene expression or knockdown**

Lentiviruses were produced using 293T cells for gene over-expression and/or knockdown, and 2 μg/ml puromycin was used for cell line selection. RNA and/or protein were collected for further gene expression analyses. Targeting sequences of *Taz*, *Brg1*, *Arid1a* and *Arid2* are listed as follows:

| Gene | shRNA#1 (5'-3') | shRNA#2 (5'-3') |
| --- | --- | --- |
| Taz | CAGCCGAATCTCGCAATGAAT | CATGAGCACAGATATGAGAT |
| Brg1 | CGCCCGACACATTATTGAGAA | TCGAGTCTCTACCAGCATTAA |
| Arid1a | CCTAGGCAGCCTAACTATAAT | TGGGCGTTAGACACCATTAAC |
| Arid2 | GACTAACAGCTGCCTTAATAT | TTCTTACTTGAGCCAATATAT |

**Western blot**

The proteins extracted from cells were used for western blot experiment. Primary antibodies were incubated at 4℃ overnight and the HRP-conjugated secondary antibodies were incubated at room temperature for 1 hr. The antibodies are listed as follows: EPCAM (Proteintech, 21050-1AP), NCAM (CST, 99746), CD44/HCAM (SantaCruz, sc-18849), TAZ (CST, 4883S), CLEAVED CASPASE 3 (CST, #9661), ASCL1 (BD pharmingen, 556604), GAPDH (ABtech, AC002), TUBULIN (proteintech, 66240-1).

**Soft agar colony formation and matrigel invasion assays**

For soft agar assay, a total of 5,000 cells in 0.4% soft agar with complete medium were seeded onto the solidified bottom layer of 1% agar. Colonies were counted after 0.005% crystal violet staining. As for matrigel invasion assay, 10,000 cells were seeded onto the solidified layer of matrigel (BD Biosciences) and grown in 2% matrigel with complete medium.

**Anoikis assay**

A total of 1×10^6^ cells were seeded in 6-well plates coated with 3% low-melting agarose for 24 hours. Cell proteins were then collected for western blot experiment.

**Reverse transcription and real-time PCR**

Total RNAs were extracted by TRIzol reagent (Life Technologies) according to the manufacturer’s instruction, and then retro-transcribed into first-strand cDNA using RevertAid™ First Strand cDNA Synthesis Kit (Fermentas). The cDNAs were then used for real-time PCR on the 7500 Fast Real-Time PCR System (Applied Biosystems) using SYBR-Green Master PCR mix (Roche). The primers sequences were as follows:

| Gene | forward primer (5'-3') | reverse primer (5'-3') |
| --- | --- | --- |
| Gapdh | ACCCAGAAGACTGTGGATGG | CACATTGGGGGTAGGAACAC |
| Ncam | ACCACCGTCACCACTAACTCT | TGGGGCAATACTGGAGGTCA |
| Cd44 | TCTGCCATCTAGCACTAAGAGC | GTCTGGGTATTGAAAGGTGTAGC |
| Taz | GAAGGTGATGAATCAGCCTCTG | GTTCTGAGTCGGGTGGTTCTG |
| Ascl1 | GCAACCGGGTCAAGTTGGT | GTCGTTGGAGTAGTTGGGGG |
| Insm1 | CCACGCCCGTGTCCTACCGG | GCAGCATGGGCGCGCTCCG |
| Neurod1 | ATGACCAAATCATACAGCGAGAG | TCTGCCTCGTGTTCCTCGT |
| Chga | ATCCTCTCTATCCTGCGACAC | GGGCTCTGGTTCTCAAACACT |
| Sox11 | CGACGACCTCATGTTCGACC | GACAGGGATAGGTTCCCCG |
| Ttf1 | CAACAACTGCAGCAGGACAG | GGGTTTGCCGTCTTTGACTA |
| Brg1 | CAAAGACAAGCATATCCTAGCCA | CACGTAGTGTGTGTTAAGGACC |
| Arid1a | TGGGACTAACCCATACTCGCA | GAATCTGCTGTGCATAAGAGAGG |
| Arid2 | ACACTGTTATCATAGCACCCCC | CAATGTTCTGGAATCCTACCCTG |
| Yap | AAGCTGCCCGACTCCTTCTTCAA | CTCCACAGCATGTTCGAGCTCA |

**The primers sequences for genotyping**

| Gene | forward primer (5'-3') |
| --- | --- |
| Brg1 k/o-1 | GCCTTGTCTCAAACTGATAAG |
| Brg1 k/o-2 | GTCATACTTATGTCATAGCC |
| Brg1 k/o-3 | GATCAGCTCATGCCCTAAGG |
| Rb1 18 | GGCGTGTGCCATCAATG |
| Rb1 212 | GAAAGGAAAGTCAGGGACATTGGG |
| Trp53-1 | CGCAATCCTTTATTCTGTTCG |
| Trp53-2 | AGCACATAGGAGGCAGAGAC |
| Trp53-3 | TGAGACAGGGTCTTGCTATTG |
